# Segregation of brain and organizer precursors is differentially regulated by Nodal signaling at blastula stage

**DOI:** 10.1101/2020.07.16.167320

**Authors:** Aitana M. Castro Colabianchi, María B. Tavella, Laura E. Boyadjián López, Marcelo Rubinstein, Lucía F. Franchini, Silvia L. López

## Abstract

The Blastula Chordin- and Noggin Expressing Center (BCNE) comprises animal-dorsal and marginal-dorsal cells of the amphibian blastula and contains the precursors of the brain and of the gastrula organizer. Previous findings suggested that the BCNE behaves as a homogeneous cell population that depends only on nuclear β-catenin activity but does not require Nodal and segregates into its descendants later, during gastrulation. In this work, we analyzed if the BCNE is already compartmentalized at the blastula stage. In contrast to previous findings, we show that the BCNE does not behave as a homogeneous cell population in response to Nodal antagonists. In fact, we found that the *chordin.1* expression in a marginal subpopulation of notochordal precursors indeed requires Nodal input. We also establish that an animal BCNE subpopulation of cells that express both, *chordin.1* and *sox2* (a marker of pluripotent neuroectodermal cells), and gives rise to most of the brain, persisted at blastula stage after blocking Nodal. Moreover, RT-qPCR analysis showed that *chordin.1* and *sox2* expression increased at blastula stage after blocking Nodal. Therefore, Nodal signaling is required to define a population of *chordin.1+* cells and to restrict the recruitment of brain precursors within the BCNE as early as at blastula stage.

## INTRODUCTION

When the dorsal lip of an amphibian early gastrula is grafted into the ventral side of a host, a secondary embryo develops, with complete anterior-posterior and dorsal-ventral axis (Spemann and Mangold, 1924), (De Robertis and Kuroda, 2004). Because of these properties, the dorsal lip region is known as the Gastrula Organizer (GO). The GO is preceded by an earlier dorsal signaling center, the “Blastula Chordin- and Noggin-Expressing Center” (BCNE), which expresses Chordin.1 (Chrd.1) and Noggin, two BMP antagonists proposed to initiate anterior neural induction directly in the BCNE (De Robertis and Kuroda, 2004). This center encompasses the animal/marginal dorsal cells of the blastula and gives rise to the forebrain and most of the midbrain and hindbrain, as well as to all endomesodermal tissues derived from the GO. Although *Chrd.1* transcripts are distributed throughout the entire BCNE, they are restricted during gastrulation to the endomesodermal descendant of this center (GO and its derivatives) but are absent from the neuroectodermal descendant (the presumptive brain) (Kuroda et al., 2004). It was proposed that dorsal accumulation of maternal nuclear β-catenin (nβ-cat) triggers *Chrd.1* expression throughout the BCNE, while Nodal signaling is only required later to maintain *Chrd.1* in the GO (Wessely et al., 2001). The siomois-related homeobox genes (sia) are directly activated by the dorsal Wnt/nβ-cat cascade. They encode the first transcription factors expressed in the BCNE and activate *Chrd.1* expression by directly binding to its promoter (Ishibashi et al., 2008), (Reid et al., 2012). Thus, it appeared that only dorsal nβ-cat signaling initiates brain and GO development through the establishment of the BCNE, while Nodal would be required later for the maintenance of the GO and its descendants. These findings suggested that the BCNE behaves as a homogeneous cell population induced by dorsal nβ-cat and segregates later, during gastrulation, into brain and GO.

Accumulation of transcripts encoding *Xenopus* Nodal-related endomesodermal inducers (Xnrs) in the Nieuwkoop Center (NC), located in the vegetal dorsal cells, requires the cooperative action of VegT and nβ-cat (Takahashi et al., 2000). Nodal activity can be experimentally blocked by the C-terminal fragment of Cerberus protein, known as Cerberus-short (Cer-S), which specifically binds to and antagonizes Xnrs (Bouwmeester et al., 1996), (Piccolo et al., 1999), (Takahashi et al., 2000). After blocking mesoderm induction with *cer-S* mRNA (hereafter, *cer-S),* embryos still develop head structures, including brain tissue and a cyclopic eye, express the pan-neural marker *sox2* and forebrain, midbrain and hindbrain markers (Wessely et al., 2001). These findings indicated that anterior neural tissue can be specified in the absence of mesoderm, lending support to the idea that anterior neural specification is initiated in the BCNE due to the transient expression of neural inducers in the presumptive brain territory (De Robertis and Kuroda, 2004).

In this work, we aimed to determine if the BCNE behaves as a homogeneous or heterogeneous cell population in response to Nodal. To this end, we inhibited the Nodal pathway with *cer-S* or with a dominant-negative form of FoxH1, a transcription factor with Forkhead domain which binds Smad2 and Smad4, the transducers of Nodal signaling (Chen et al., 1997), (Watanabe and Whitman, 1999), (Hill, 2001). We found that the BCNE is functionally and topologically compartmentalized, as revealed by a differential response of distinct cell subpopulations to the blockade of Nodal. We demonstrate that Nodal is already necessary as early as at blastula stage for restricting a subpopulation of brain precursors while favoring a subpopulation of chordal organizer precursors. We also found that, during gastrulation, Nodal is required for maintaining the axial mesoderm (AM), with the chordal mesoderm (CM) being the subpopulation with highest sensitivity to Nodal depletion. Finally, we compare the requirement of Nodal signaling in the segregation of dorsal territories in *Xenopus* with those previously observed in other vertebrate models.

## RESULTS

### BCNE cells do not respond uniformly to Nodal blockade

We blocked Nodal by injecting *cer-S* and analyzed at s9 by *in situ* hybridization (ISH) if this could result in spatial changes of the BCNE marker *Chrd.1* and its up-stream regulator, the direct Wnt/nβ-cat target gene *sia1* (Ishibashi et al., 2008), (Reid et al., 2012). Notably, *Chrd.1* expression decreased in the marginal region of the BCNE (red asterisk, Fig. 1B) in comparison to control siblings (Fig. 1A), remaining intact in the animal region (green asterisk, Fig. 1B) (Table 1). This suggests that the BCNE is composed of two *Chrd.1* subdomains, regarding their response to Nodal blockade. However, the domain of the up-stream regulator *sia1* was not reduced after *cer-S* injection at any place at s9 (Fig. 1D,E; Table 1). These results indicate that Nodal is required for *Chrd.1* expression in a subpopulation of BCNE cells, regardless of *sia1.*

**Figure 1.**
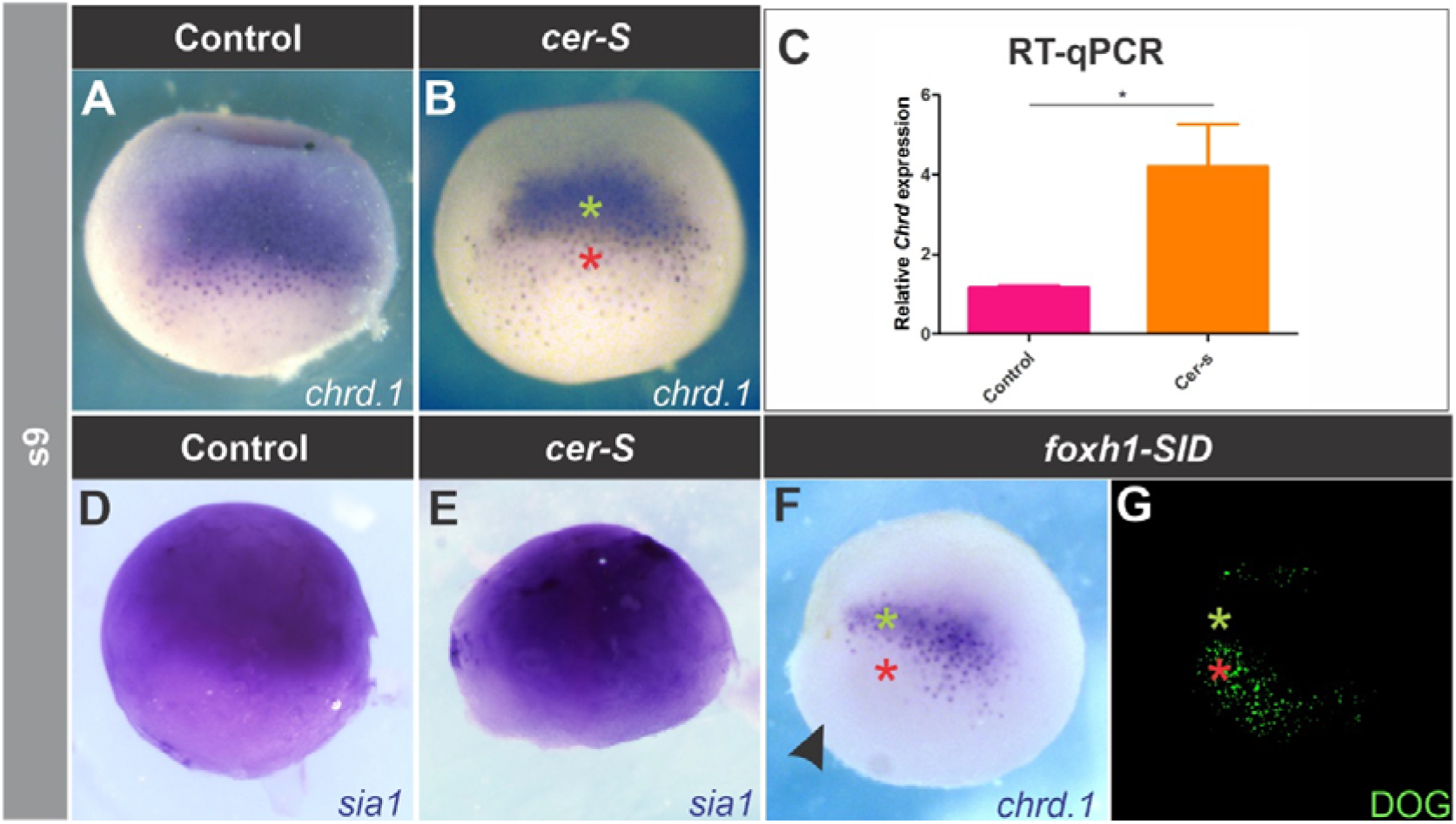
Effects of blocking Nodal on *Chrd.1* (A-C, F,G) and *sia1* expression (D,E) at late blastula (s9). *Chrd.1* (A) and *sia1* (D) are normally expressed in the whole BCNE center. *Cer-S* (B) and *foxh1-SID* (F,G) injections revealed a marginal BCNE subpopulation of cells that depends on Nodal to express the neural inducer *Chrd.1* (red asterisk), while its up-stream regulator *sia1* (E), a direct Wnt/nβ-cat target, and the animal *chrdl+* subdomain in the BCNE (B,F, green asterisk), do not depend on Nodal. The *foxh1-SID-injected* side is evidenced by DOG fluorescence (G) and is indicated by a black arrowhead. (C) RT-qPCR analysis showed a significant increase (p<0.05) in the levels of *Chrd.1* transcripts as a result of *cer-S* injection (p=0.027, unpaired, two-tailed t-test) when compared with uninjected siblings. Bars represent mean + S.E.M. of 6 biological replicates.

**Table 1.**
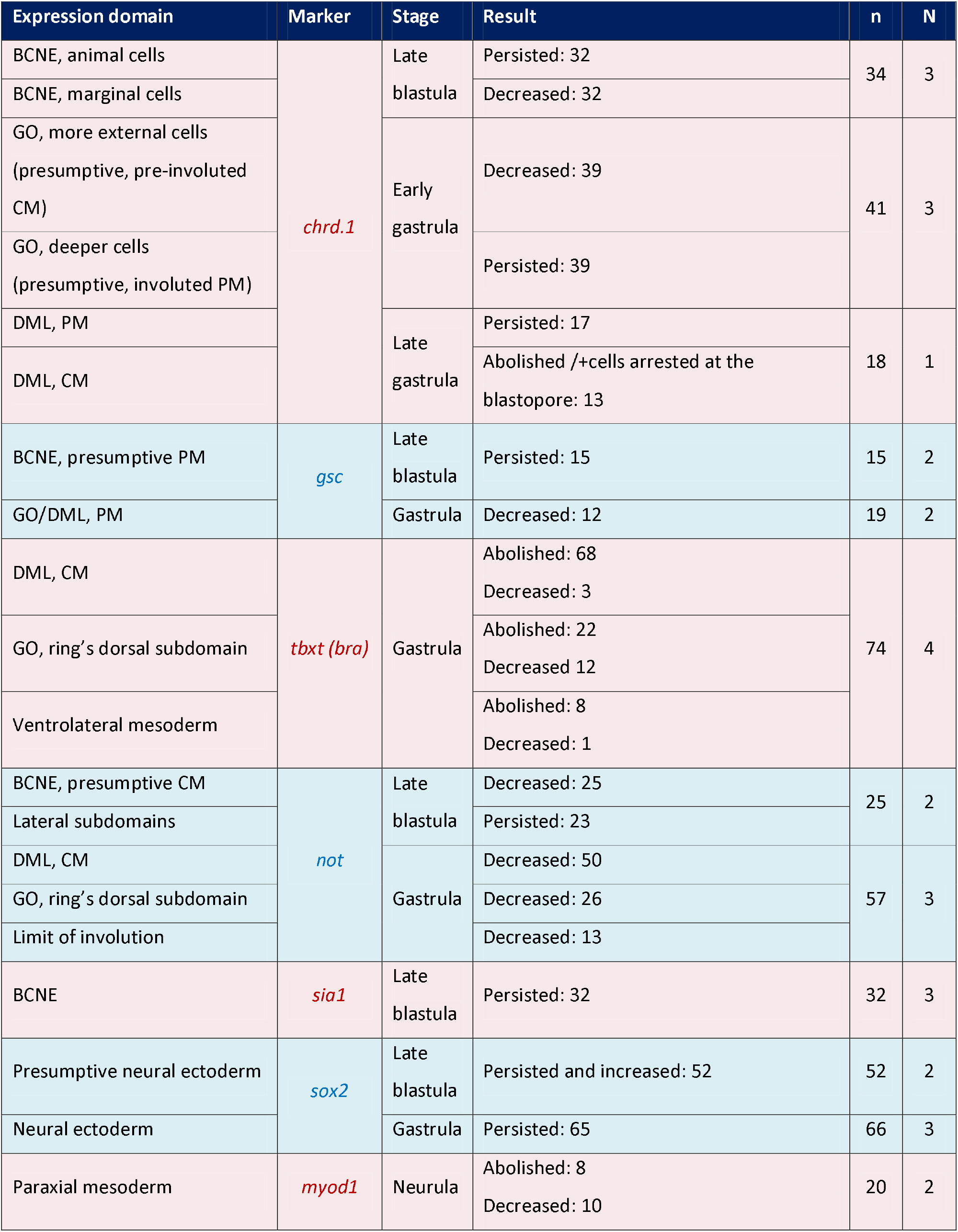
Effects of cer-S on markers of BCNE and its derivatives (GO, DML, neural ectoderm), pan-mesoderm *(tbxt/bra),* and paraxial mesoderm *(myod1).* Spatial expression was analyzed by ISH. Results are expressed as the total number of embryos showing the indicated effect on each marker. DML, dorsal midline; n, number of injected embryos; N, number of independent biological replicates.

To corroborate this, we unilaterally injected *foxh1-SID* mRNA (hereafter, *foxh1-SID),* which encodes a dominant inhibitor of Smad2 activity and blocks Nodal signaling in a cell-autonomous manner (Chen et al., 1997), (Müller et al., 2000). By comparing the injected- and non-injected sides at s9, we found that the animal expression of *Chrd.1* was not affected (green asterisk, Fig. 1F,G) but was suppressed in the marginal subdomain of the BCNE (red asterisk, Fig. 1F,G; Table 2).

**Table 2.**
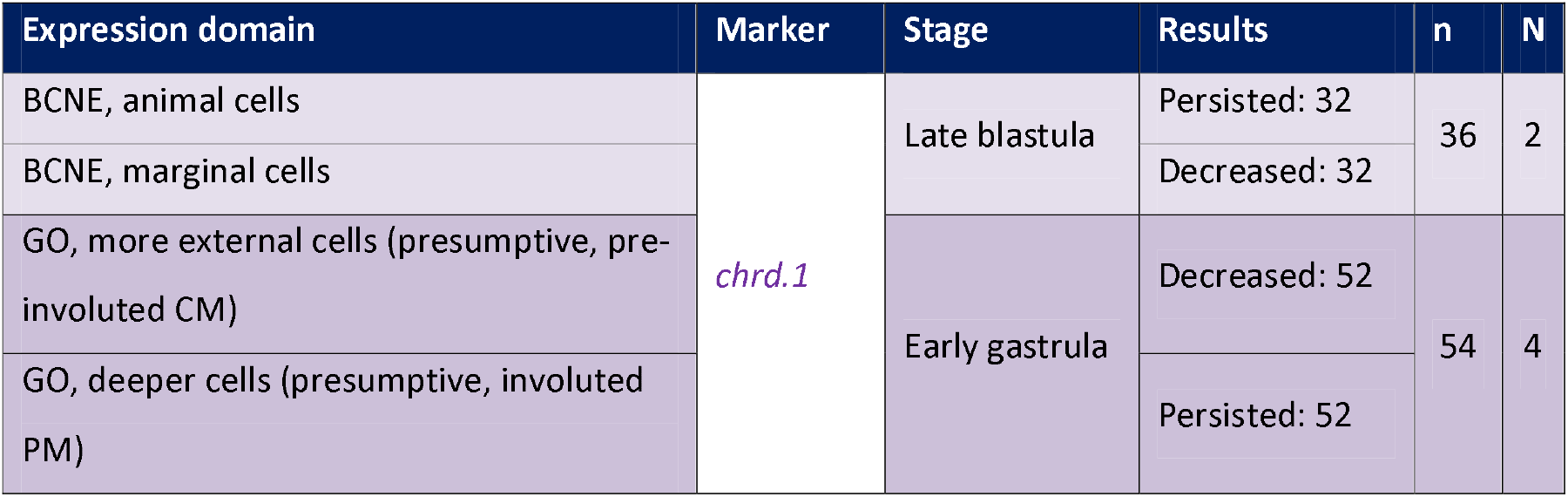
Effects of *foxh1-SID* on the spatial expression of *Chrd.1* in the BCNE and in the GO. Expression was analyzed by ISH. Results are expressed as the total number of embryos showing the indicated effect on *Chrd.1.* n, number of injected embryos; N, number of independent biological replicates.

These results confirm that the BCNE is already compartmentalized into two *Chrd.1* subdomains: 1) an animal subdomain, which persists after blocking Nodal, and 2) a marginal subdomain, which depends on Nodal. This finding is in contrast to a previous proposal which stated that Nodal is not necessary for *Chrd.1* expression in the BCNE and it is only required later for its maintenance in the GO (Wessely et al., 2001).

### Mesodermal derivatives of the BCNE display a differential response to Nodal blockade

We next analyzed the consequences of blocking Nodal on *Chrd.1* expression during gastrulation, when it is not expressed anymore in brain precursors but persists in the GO and in its AM descendants. At early gastrula, the presumptive prechordal mesoderm (PM), which is the first to involute, occupies a relatively more internal position than the pre-involuted, presumptive CM, and both express *Chrd.1* (Zorn et al., 1999), (Kaneda and Motoki, 2012). At this stage, we found a less dense ISH staining for *Chrd.1* in cer-S-injected embryos in comparison to uninjected control siblings (Fig. 2A-C). *Chrd.1* was downregulated or suppressed from more external GO cells, which correspond to pre-involuted prospective CM at the beginning of gastrulation. As these cells became more transparent due to the suppression of *Chrd.1* staining, we could observe that *Chrd.1* expression persisted in a cloud below them (Fig. 2B,C; Table 1). This deeper expression, more refractory to *cer-S,* correspond to the presumptive, involuted PM at this stage. Unilateral injection of *foxh1-SID* (0.25 ng to 1 ng) allowed to better compare the behavior of these subpopulations between the injected and the non-injected sides. *Chrd.1* persisted (green asterisk, Fig. 2G,I) or the domain was even expanded (light blue asterisk, Fig. 2H) in the deeper, involuted cells (Table 2), while in the preinvoluted, more external cells, *Chrd.1* was suppressed (red asterisk, Fig. 2G-I; Table 2). To corroborate the contribution of *chrd+* cells to PM and CM, we analyzed the expression of this marker at s13. Normally, at this stage, all *chrd+* cells have been internalized, and PM and CM have completely segregated. This allows the distinction of two *chrd+* subdomains in the AM, with clear distinct shapes: a) an anterior fan-like subdomain, characterized by an active migration behavior, corresponding to the PM (green arrow, Fig. 2D); b) a posterior subdomain, corresponding to the notochordal cells (red arrow, Fig. 2D), which ultimately form a rod by convergent-extension movements (Murgan et al., 2014), (Kwan and Kirschner, 2003). *Cer-S-* injected embryos showed that most *Chrd.1* expression was comprised by the fan-shaped subdomain (green arrow, Fig. 2E,F). In contrast, the posterior subdomain was lost or reduced (red arrows, Fig. 2E,F), sometimes with *Chrd.1+* cells arrested on the blastopore, unable to involute (yellow asterisk, Fig. 2E) (Table 1).

**Figure 2.**
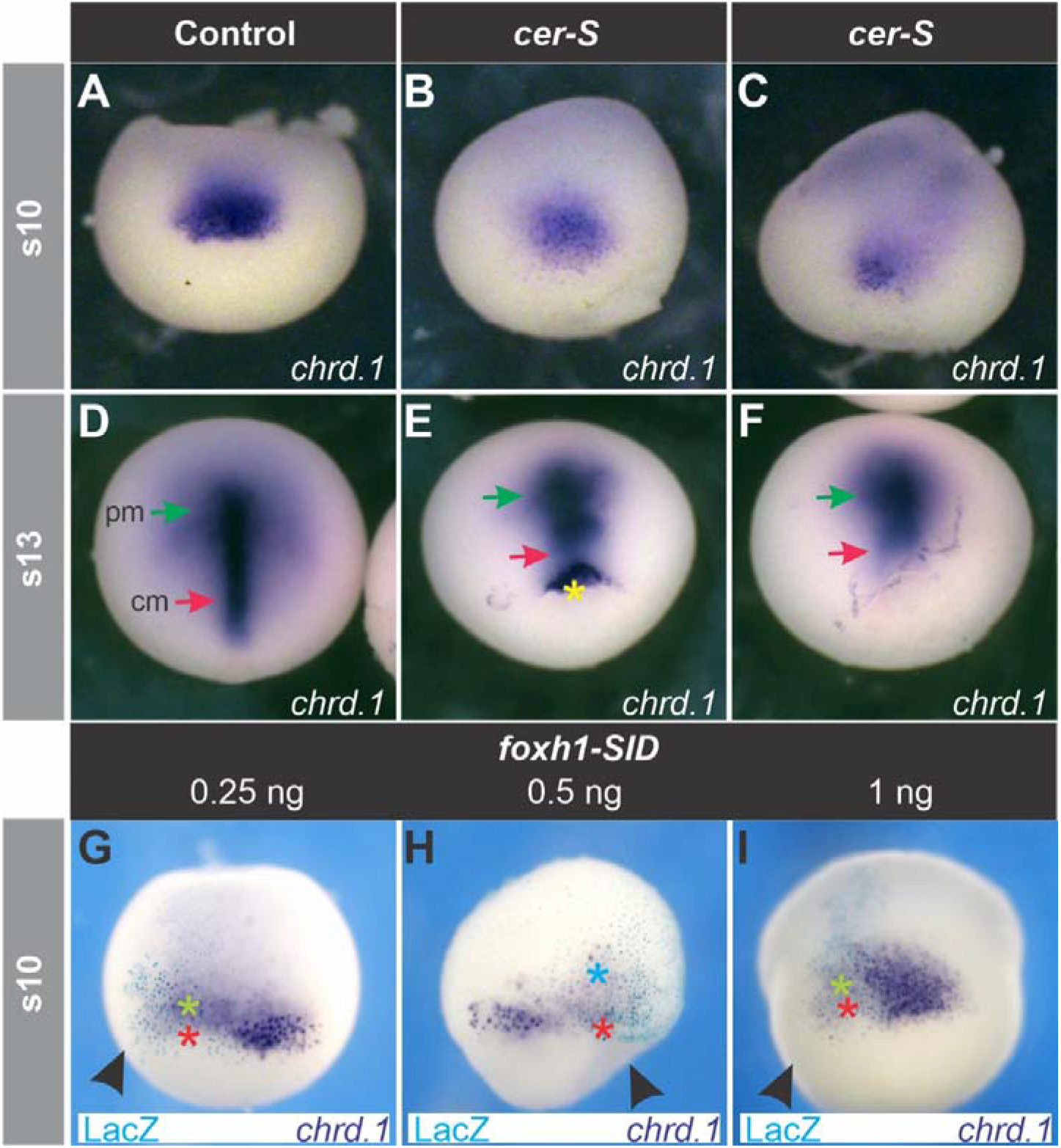
Effects of blocking Nodal on *Chrd.1* expression during gastrulation. (B,C,E,F) Injection of *cer-S.* (G-I) Unilateral injection of *foxh1-SID.* (A,D) Uninjected control siblings of embryos shown in (B,C) and (E,F), respectively. At early gastrula (s10), *Chrd.1* is expressed in the more external, pre-involuted presumptive CM cells and in the deeper, involuted presumptive PM, which are seen together as a compact domain (A, and non-injected side in G-I), but ceases expression in brain precursors (Kuroda et al., 2004). *Cer-S* (B,C) and *foxh1-SID* (G-I) suppressed *Chrd.1* expression in the pre-involuted population (red asterisks, *G-I*), but did not affect (green asterisks, G-I) or even expanded (light blue asterisk, H) *Chrd.1+* cells in the involuted population. The *foxh1-SID-injected* side is evidenced by Xgal turquoise staining (G-l) and is indicated by black arrowheads. At late gastrula (s13), *Chrd.1* is expressed in the PM (green arrow, D) and in the CM (red arrow, D). In cer-S-injected embryos, PM *Chrd.1* expression persisted (E,F, green arrows) but CM expression decreased (E,F, red arrows) or was arrested at the blastopore (E, yellow asterisk). All embryos are shown in dorsal views, anterior side upwards.

It was previously shown that not all the PM express *Chrd.1* in *Xenopus* (Yamaguti et al., 2005). These authors showed that the PM consists of two cell populations, a more anterior one (APM) expressing the homeodomain transcription factor Goosecoid (Gsc), and a more posterior one (PPM) expressing *Chrd.1* (Yamaguti et al., 2005). The same occurs in mouse (Anderson et al., 2002). Therefore, we analyzed the consequences of *cer-S* injection on the spatial expression of *gsc* and of the CM-specific transcription factor Tbxt/Brachyury, which control the characteristic movements of PM and CM cells, respectively, in an antagonistic way (Artinger et al., 1997), (Latinkić and Smith, 1999), (Kwan and Kirschner, 2003), (Luu et al., 2008). Our ISH results show that, at gastrula stages, *gsc* expression decreased in most cer-S-injected embryos (Fig. 3F,G) (Table 1). We also found through RT-qPCR analysis that *gsc* was down-regulated at the onset of gastrulation after overexpression of *cer-S* (Fig. 3H). ISH analysis showed that *tbxt* was affected at different degrees. The expression normally found in the extending notochord (red arrows, Fig. 4A,C,E) was suppressed in almost all injected embryos (Fig. 4B,D,F,G; Table 1). The expression in the GO presumptive mesoderm (dorsal part of the *tbxt* ring, yellow asterisk, Fig. 4A,C,E) often disappeared or was down-regulated (Fig. 4B,D,F; Table 1) and that of the non-GO presumptive mesoderm (white arrows, Fig. 4A,C,E) was sometimes suppressed (Fig. 4B,G; Table 1). As an independent notochordal marker, we also analyzed the spatial expression of *not,* which encodes a homeodomain transcription factor normally expressed during gastrulation in the extending CM, GO, and in the limit of involution (von Dassow et al., 1993). *Cer-S* injections affected the *not* pattern in a similar way to *tbxt* (Fig. 4H-M; Table 1). These observations indicate that, during gastrulation, among the mesodermal descendants, CM cells are the most sensitive to Nodal blockade, which disrupted notochord development almost completely. Although cells with PM characteristics persisted, as shown by ISH of *Chrd.1,* a lower *gsc* expression was detected at gastrula stages, both by ISH and by RT-qPCR. This might be due to a differential sensitivity among APM and PPM subpopulations to Nodal blockade. In addition, the ventrolateral mesoderm was less sensitive to *cer-S* than the CM, as shown by a lower penetrance of the downregulation of tbxt in the circumblastoporal ring.

**Figure 3.**
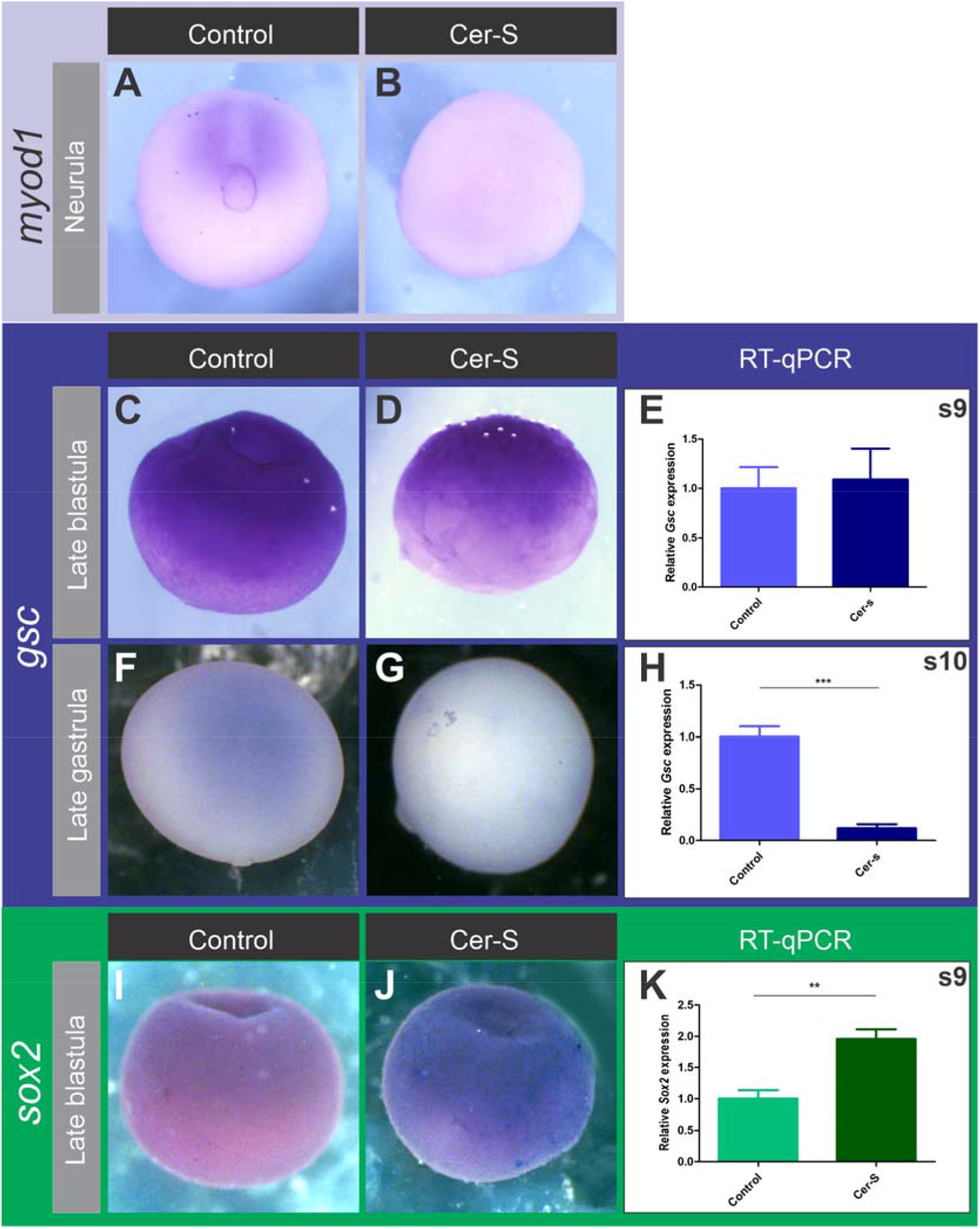
Effects of *cer-S* on the paraxial mesoderm marker *myod1* (A,B), the PM marker *gsc* (C-H), the neural marker *sox2* (I-K) at the stages indicated. (A) Control neurula showing *myod1* expression, which was abolished or drastically reduced in cer-S-injected siblings (B). (C-E) After *cer-S* injection, *gsc* expression was not significantly affected at s9, as revealed by ISH (D) and RT-qPCR (E) (p=0.8184; unpaired, two-tailed t-test) when compared with uninjected siblings. Bars represent mean + S.E.M. of 6 biological replicates. (F-G) At the end of gastrulation, *gsc* expression decreased in the PM of most cer-S-embryos injected, as revealed by ISH (G). *Gsc* transcripts levels were also significantly reduced at s10 (p<0.05), as revealed by RT-qPCR (H) (p=0.0002; unpaired, two-tailed t-test) when compared with uninjected siblings. Bars represent mean + S.E.M. of 4 biological replicates. (I) Control blastula showing expression of the neural marker *sox2,* which was consistently increased in cer-S-injected siblings, as revealed by ISH (J). A significant increase (p<0.05) in *sox2* transcripts levels was detected by RT-qPCR at s9 in cer-S-injected embryos (p=0.0016; unpaired, two-tailed t-test) when compared with uninjected siblings. Bars represent mean + S.E.M. of 5 biological replicates. (A,B), posteriordorsal views; (C,D,F,G,I,J) Dorsal views.

**Figure 4.**
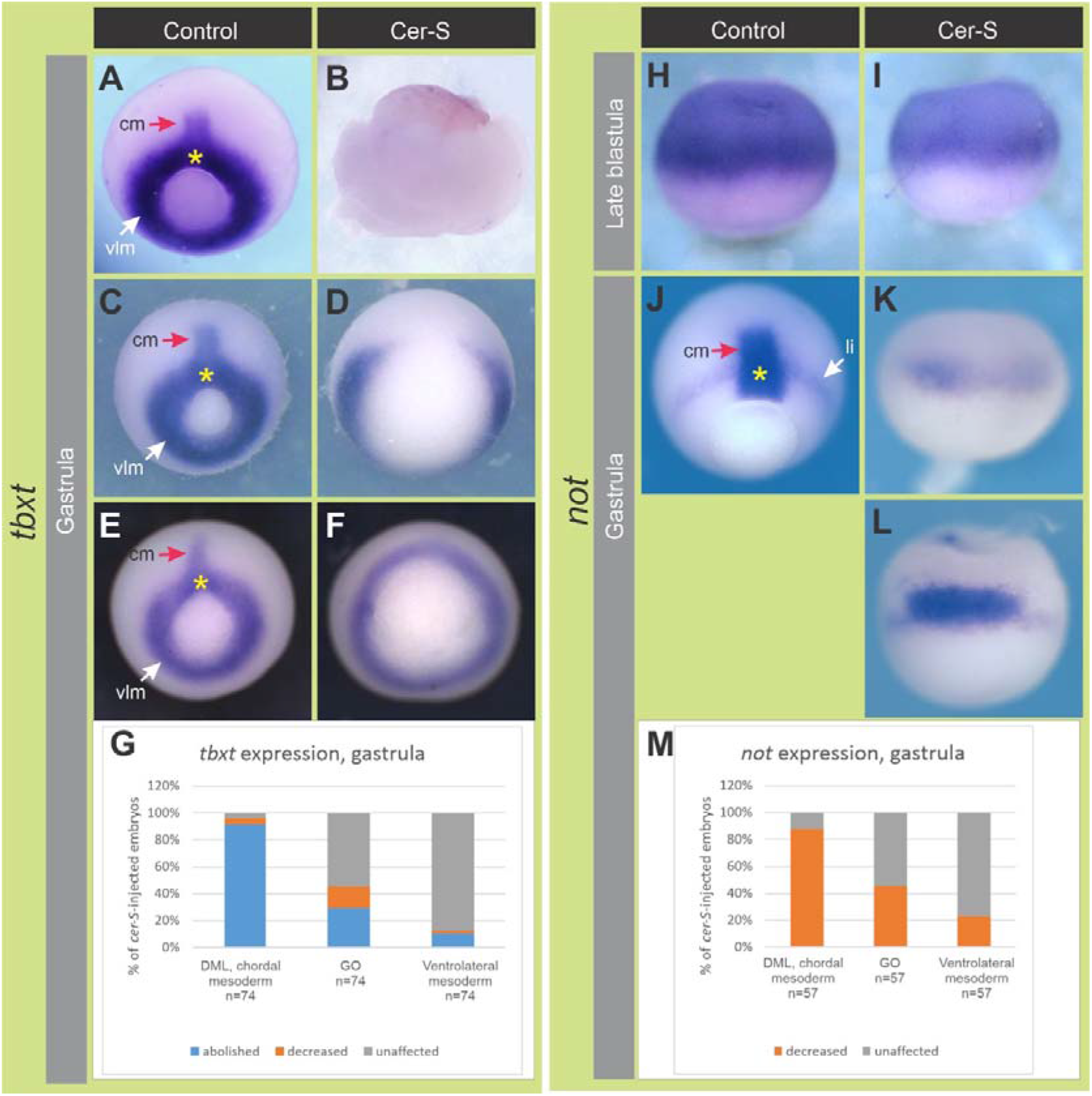
Effects of *cer-S* on the spatial expression of the pan-mesoderm/CM marker *tbxt* (AG) and the CM marker *not* (H-M) at the stages indicated in grey boxes. (A,C,E) Control gastrulae showing *tbxt* expression in the CM (red arrow), GO (yellow arrow) and presumptive ventrolateral mesoderm in the blastopore (vlm, white arrow). (B,D,F) cer-S-injected embryos which are siblings of those shown in A,C,E, respectively. (G) Summary of the effects of *cer-S* on *tbxt* expression at gastrula stage. Results are expressed as percentage of embryos showing the indicated phenotypes for each *tbxt* subdomain. (H) Dorsal view of a control late blastula, showing strong *not* expression in the BCNE region, which decreased in cer-S-injected siblings (I). (J) Control gastrula showing *not* expression in the extending CM (red arrow), GO (yellow asterisk) and limit of involution (li, white arrow). *Cer-S* abolished *not* expression in the extending notochord (K,L,M) and often decreased it in the GO (K,M); when *not* expression was not decreased in the GO (L), *not+* cells were arrested at the blastopore, unable to involute. (M) summary of the effects of *cer-S* on *not* expression at gastrula stage. Results are expressed as percentage of embryos showing the indicated phenotypes for each *not* subdomain. See main text for details. (A-F,J), posterior/dorsal views. (H,I,K,L) Dorsal views.

In conclusion, Nodal is required for the onset of *Chrd.1* expression in the marginal subdomain of the BCNE at blastula stage. Later, the mesodermal descendants of the BCNE require Nodal for their maintenance during gastrulation, with the CM having the highest requirement.

### Nodal restricts neural specification and is required for the expression of CM but not of PM markers at the BCNE stage

We have observed that blocking Nodal distinguishes two *Chrd.1+* subpopulations in the BCNE. Since this center is composed of brain and GO precursors, and the GO, in turn, segregates into PM and CM, we wondered if blocking Nodal could already discriminate between PM and CM precursors and the neural ectoderm at s9. For this purpose, we compared the expression of the pan-neural marker *sox2* and the specific PM (gsc) and CM *(not)* markers in the BCNE between cer-S-injected embryos and uninjected control siblings at s9. Although it is known that *gsc* and *tbxt* are involved in an antagonistic relationship that controls PM and CM segregation during gastrulation (Artinger et al., 1997), (Latinkić and Smith, 1999), (Kwan and Kirschner, 2003), (Luu et al., 2008), *tbxt* is weakly expressed in the presumptive CM at late blastula. Therefore, we chose *not* as an alternative spatial marker of CM precursors. While *gsc* and *sox2* expression persisted and even increased in the case of the latter (Fig. 3C-E, I-K; Table 1), *not* expression in the BCNE subdomain corresponding to the presumptive CM territory decreased in cer-S-injected embryos (Fig. 2H,I; Table 1). Interestingly, RT-qPCR analysis showed that *Chrd.1* expression at s9 increased after blocking Nodal (Fig. 1C) (see Discussion). Thus, we propose that BCNE cells are functionally compartmentalized, with presumptive CM cells requiring Nodal for their specification, while neural specification is restricted by Nodal.

## DISCUSSION

The results presented here indicate that Nodal is indeed required to trigger the full expression of the neural inducer *Chrd.1* in the BCNE (Fig. 5C), in contrast to a previous proposal (Wessely et al., 2001) (Fig. 5A,B). In fact, we found that *Chrd.1* was abolished by either *cer-S* or *foxh1-SID* in the marginal region of this center. In contrast, *Chrd.1* persisted in the animal subdomain of the BCNE after blocking Nodal, as revealed by ISH. Furthermore, RT-qPCR analysis showed that both, *Chrd.1* and *sox2* expression increased at s9 in cer-S-injected embryos. These results indicate that the population of brain precursors expressing both, the neural inducer and the neural specification marker in the BCNE, was expanded after blocking Nodal. Indeed, we notice that the *Chrd.1* domain in cer-S-injected embryos at s9 shown in Fig. 3D,E in (Wessely et al., 2001) is less extended in the animal-vegetal axis in comparison to control siblings, resembling the results shown in the present work, but the authors interpreted that *Chrd.1* expression was refractory to the blockade of Nodal. In addition, we found that the PM marker *gsc* persisted whereas the notochordal marker not was reduced in the BCNE after blocking Nodal. Therefore, mesodermal precursors in the BCNE are differentially regulated by Nodal.

**Figure 5.**
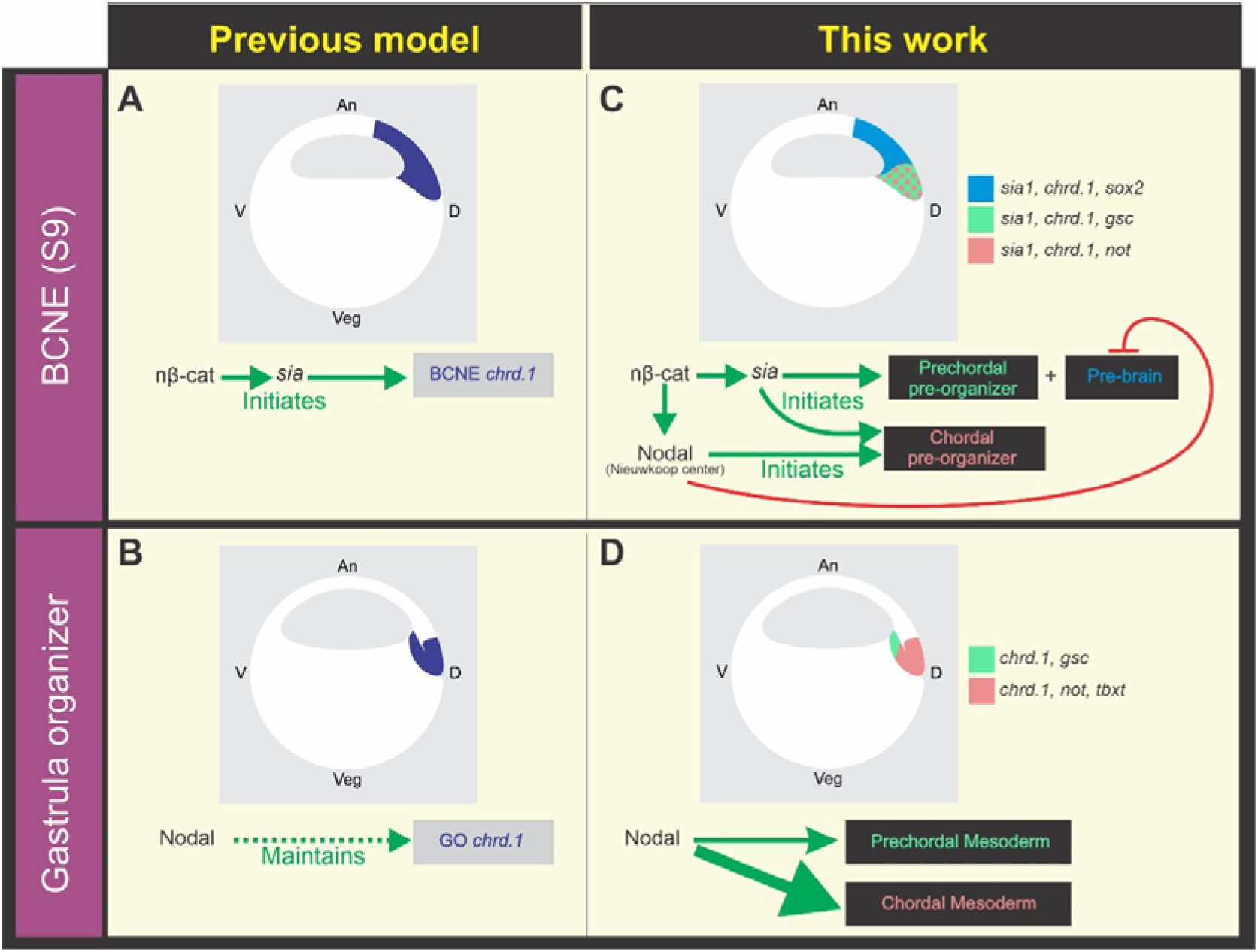
(A,B) Previous model of Wnt/nβ-cat and Nodal requirements for *Chrd.1* expression in the blastula and gastrula dorsal signaling centers (Wessely et al., 2001). Dorsal nβ-cat initiates *Chrd.1* expression in the BCNE through direct activation of the gene encoding the transcription factor Sia (Ishibashi et al., 2008). According to this model, Nodal signaling is not required to initiate *Chrd.1* expression in the BCNE (A), but it is later required for the maintenance of *Chrd.1* expression in the GO (B). (C,D) Role of Nodal and Wnt/nβ-cat in BCNE compartmentalization and in the development of its derivatives updated in the present work. The expression domains of the markers analyzed in this study are color-coded. (C) Dorsal nβ-cat initiates pre-brain and prechordal pre-organizer induction through the activation of *sia* in the BCNE. Accumulation of *nodal* transcripts in the NC requires the cooperative action of VegT and dorsal nβ-cat (Takahashi et al., 2000). Dorsal nβ-cat initiates chordal pre-organizer induction through the activation of *sia* in the BCNE and *Xnrs* in the NC. (D) During gastrulation, high Nodal signaling maintains CM development, whereas low Nodal signaling maintains PM development.

Altogether, these results indicate that the BCNE does not behave as a homogeneous cell population. In fact, our data show that the BCNE is compartmentalized into the precursors of the prospective brain (pre-brain), the PM precursors and the CM precursors, which can be distinguished earlier than previously thought, since they are being differentially regulated by Nodal (Fig. 5C). Therefore, we present here a modified model of the originally proposed by Wessely et al. (2001), concluding that *Chrd.1* expression is triggered in the whole BCNE by Wnt/nβ-cat through *sia* (Ishibashi et al., 2008) and has different requirements for Nodal, depending on the BCNE sub-population. Nodal is necessary to trigger *Chrd.1* expression in the chordal pre-organizer, whereas it is not required at the prechordal pre-organizer. At the prebrain subpopulation, *Chrd.1* is restricted by Nodal (Fig. 5C). Since *sia1* expression persisted in the BCNE after *cer-S* injection and *sox2* expression increased, we suggest that Nodal restricts the pre-brain territory downstream of *sia* (Fig. 5C). In addition, within the presumptive mesodermal subdomain in the BCNE (pre-organizer), Nodal promotes the development of posterior AM derivatives but does not affect the presumptive prechordal subpopulation at this stage (Fig. 5C). Later, Nodal is necessary to maintain both the CM and the PM during gastrulation. The *Chrd.1, tbxt* and *gsc* patterns at gastrula stages suggest that, among all mesodermal derivatives, the CM is the most sensitive to the blockade of Nodal in *Xenopus.* Thus, it appears that CM maintenance during gastrulation requires more input from Nodal than other mesodermal cell types, like the PM.

The default-state model of neural specification adopted for *Xenopus* and mouse embryos poses that the default fate of ectoderm (epiblast) is neural, but it is actively repressed by BMP signaling. This default-state is revealed during neural induction, when the ectoderm is exposed to BMP antagonists, like Chordin and Noggin (Levine and Brivanlou, 2007). In *Xenopus,* this initial step of neural induction occurs before gastrulation in the BCNE, as the BMP antagonists are directly expressed by neuroectodermal precursors fated to give rise to the forebrain (Kuroda et al., 2004). Like in *Xenopus* at blastula stage (this work, Fig. 6A), the first neural tissue induced in mouse expresses the pan-neural marker Sox2 (Levine and Brivanlou, 2007). Moreover, this neural tissue initially has a forebrain character, but it is subsequently posteriorized during gastrulation to form the remainder of the central nervous system (CNS) (Levine and Brivanlou, 2007) (Fig. 6I,J). However, unlike *Xenopus,* Chrd and Noggin are not expressed before gastrulation in mouse. Chrd transcripts first appear in the GO at mid-streak stage (Anderson et al., 2002), (Kinder et al., 2001) (Fig. 6K). Noggin appears later in the GO, and both genes are expressed in the GO-derived AM (Fig. 6L) (McMahon et al., 1998), (Bachiller et al., 2000), but they were not detected in the prospective forebrain, unlike in the *Xenopus* BCNE (Fig. 6A-C) (Kuroda et al., 2004). Therefore, the default model for anterior neural induction in mouse implied that the source of BMP antagonists lies in the GO and its derived tissues (Levine and Brivanlou, 2007), i.e., outside the presumptive forebrain. However, we note that transcripts from a Chordin-like1 gene (Chrdl1) are first detected in the future neural plate at E7.0 in mouse (Fig. 6K) and persist there during gastrulation (Fig. 6L) (Coffinier et al., 2001). Notably, Chrdl1 never appears in the node or in the primitive streak (PS), thus establishing a complementary pattern in relation to Chrd1 (Fig. 6K,L) (Coffinier et al., 2001). Chrdl1 behaves as a BMP antagonist (Sakuta et al., 2001), (Chandra et al., 2006) and mouse Chrdl1 was more potent than *Xenopus Chrd.1* in the induction of complete secondary axes in frogs (Coffinier et al., 2001). Thus, as early as at E7.0 [when a range of pre-streak to mid-streak embryos can be obtained, according to EMAP eMouse Atlas Project (http://www.emouseatlas.org)] the mouse future neural plate indeed expresses a BMP antagonist of the Chordin family. An ortholog *chrdl1* gene was identified in *Xenopus,* but transcripts were not detected before tailbud stages (Pfirrmann et al., 2014). Therefore, it seems that cells in the presumptive brain territory already express BMP antagonists as early as at late blastula in *Xenopus (Chrd.1)* or around the onset of gastrulation in mouse (Chrdl1), when the anterior AM just begins migrating from the early GO to completely underlie the future forebrain later in gastrulation.

**Fig. 6.**
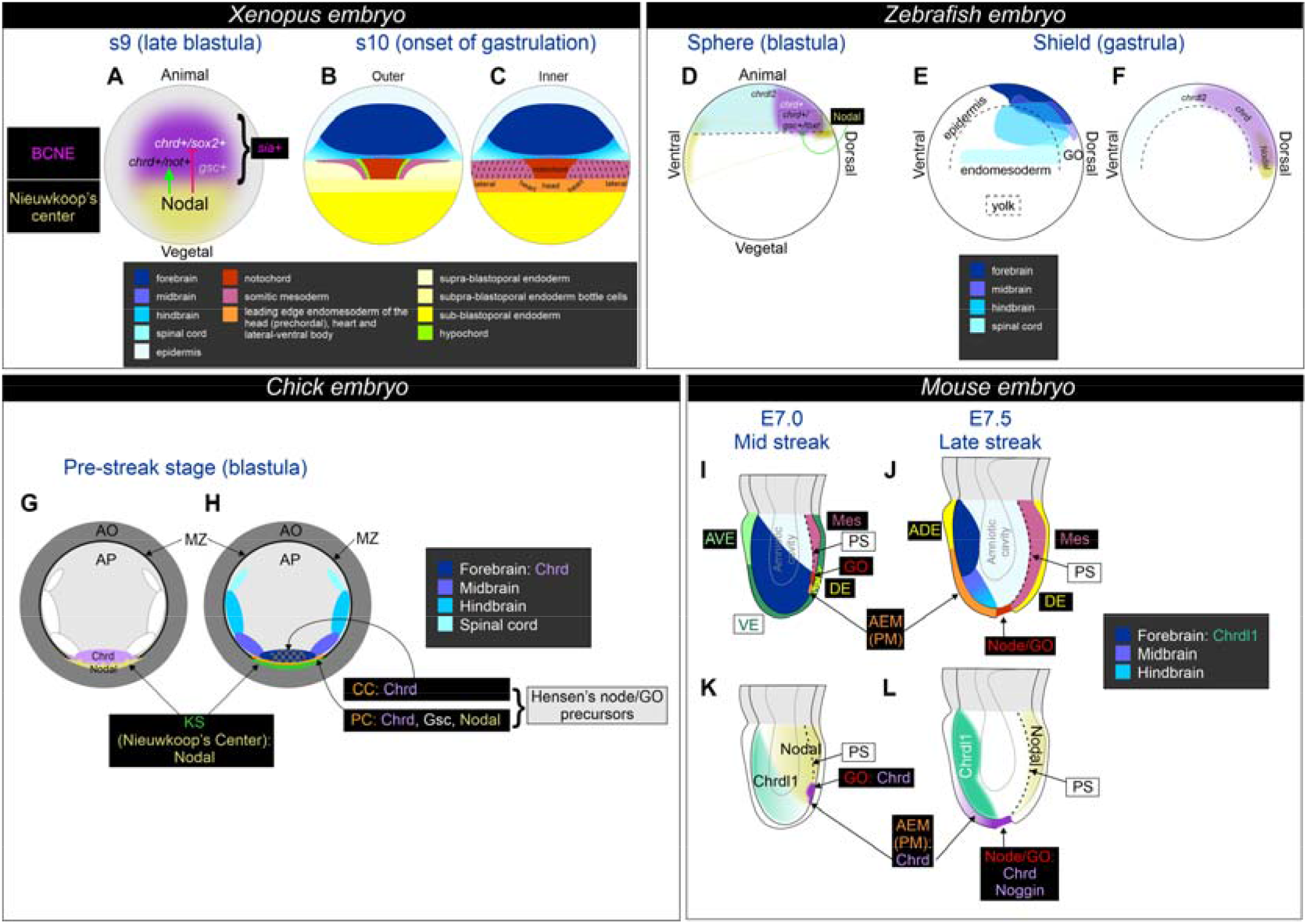
Comparison between vertebrate models. (A) Position of the dorsal signaling centers at blastula stage in *Xenopus* (dorsal view). Expression patterns were obtained from the following sources: *Chrd.1,* (Kuroda et al., 2004); *gsc,* (Sudou et al., 2012); *nodal(s),* (Agius et al., 2000), (Takahashi et al., 2000), (Kuroda et al., 2004), (Reid et al., 2016); *sia,* (Sudou et al., 2012); *sox2* (this work). While *sia* is expressed in the whole BCNE, *gsc* transcripts are present in a subset of BCNE cells (Sudou et al., 2012) and its expression is not perturbed by blocking Nodal (this work). The presence of different cell subpopulations in the BCNE are shown according to the response of the indicated markers to Nodal. The green arrow denotes that Nodal favors the specification of subpopulations expressing *Chrd.1* and *not* (black letters) and does not necessarily imply direct regulation of these genes. The red broken line denotes that Nodal restricts the specification of the subpopulations expressing *chrd* and *sox2* (white letters) and does not necessarily imply direct regulation of these genes. (B, C) Fate map of the outer (A) and inner (B) cell layers of *Xenopus* at the onset of gastrulation (dorsal view), just before the beginning of endomesoderm internalization [adapted from (Keller, 1975), (Keller, 1976), (Shook and Keller, 2004), (Shook et al., 2004)]. A high-resolution fate map of s9 *Xenopus* embryos is not available, but lineage tracing experiments demonstrated that the BCNE gives rise to the forebrain and part of the midbrain and hindbrain (neurectoderm derivatives) and to the AM and floor plate (GO derivatives) (Kuroda et al., 2004). Therefore, a rough correlation of the predicted territories can be projected from the s10 map to the s9 embryo. (D) Diagram of a sphere stage zebrafish embryo, showing the expression patterns of the following markers: *chrd* (Sidi et al., 2003), (Branam et al., 2010); *chrdl2* (Branam et al., 2010); *nodal1 (squint) + nodal2 (cyclops)* (Feldman et al., 1998), (Rebagliati et al., 1998). The presence of different cell subpopulations in the blastula dorsal signaling center are shown according to the response of the indicated markers to Nodal. The green arrow denotes that Nodal favors the specification of subpopulations expressing *chrd* and *gsc* (black letters) and does not necessarily imply direct regulation of these genes. Another subpopulation of *chrd+* cells (white letters) does not require Nodal at blastula stage. (E) Fate map for the zebrafish CNS at the shield stage. (F) *Chrd, chrdl2* and *Nodal* expression in zebrafish at shield stage (bibliographic references as in D). The broken line depicts the limit of the yolk cell. (G,H) Diagram illustrating a pre-streak stage chick embryo, showing Chrd and Nodal expression (G) and a rough fate map for the precursors of the CNS and GO (H). Predictions of the locations of the centers of the prospective territories of the CNS are shown in a gradient of blue colors, as there is great overlap of cell fates at this stage [modified from (Stern et al., 2006), (Foley et al., 2000)]. The posterior cells (PC) and central cells (CC) contributing to Hensen’s node are also shown [modified from (Streit et al., 2000)]. Expression patterns were obtained from the following sources: Chrd, (Streit et al., 1998), (Matsui et al., 2008); Nodal, (Matsui et al., 2008); Gsc, (Izpisúa-Belmonte et al., 1993). Notice the proposed overlap in the territories of the prospective forebrain (dark blue) and of a subset of the GO precursors (stippled orange), both expressing Chrd, which is also expressed by another population of GO precursors (PC) that also expresses Gsc. AO, area opaca; AP, area pellucida; KS, Köller’s sickle (located in the superficial part of the posterior marginal zone); GO, Hensen’s node; MZ, marginal zone. (I, J) Gastrulating mouse embryos at mid (I) and late (J) streak stages, respectively, indicating signaling centers and embryological regions. Illustrations were based on the following sources: (Tam and Behringer, 1997), (Tam et al., 1997), (Beddington and Robertson, 1999), (Kinder et al., 2001), (Yamaguchi, 2001), (Tam and Rossant, 2003), (Levine and Brivanlou, 2007), (Shen, 2007). ADE, anterior definitive endoderm; AEM, anterior endomesoderm; AVE, anterior visceral endoderm; GO, gastrula organizer; Mes, mesoderm; PM, prechordal mesoderm; PS, primitive streak (broken line); VE, visceral endoderm. Blue colors represent the progressive anterior-posterior regionalization of the neural ectoderm in the model for anterior neural induction/posteriorization proposed by (Levine and Brivanlou, 2007). (K,L) Expression patterns at mid (K) and late (L) streak stages were obtained from the following sources: Chrd, (Bachiller et al., 2000); Chrdl1, (Coffinier et al., 2001); Nodal, (Varlet et al., 1997), (Shen, 2007).

Single knock-out mice for Chrd or Noggin undergo normal neural induction and anterior CNS patterning (Bachiller et al., 2000), (McMahon et al., 1998). Although double knock-outs demonstrated that both are necessary for forebrain maintenance (Bachiller et al., 2000), it was not studied if forebrain specification, which normally occurs around the mid-streak stage (Levine and Brivanlou, 2007), occurred. On the other hand, Chrdl1^-/-^ mice developed a CNS and survived to adulthood (Blanco-Suarez et al., 2018). Double knockouts of Chrd and Chrdl1 should be obtained to study if both genes are necessary for neural specification.

In *Xenopus, Chrd.1* expression in the chordal pre-organizer population requires induction by Nodal (this work, Fig. 6A). We notice that in the mid-streak stage mouse, while Nodal is strongly expressed in the posterior-proximal quadrant of the epiblast, where the PS forms and endomesoderm ingression takes place (Varlet et al., 1997) (Fig 6K), Chrdl1 is oppositely expressed with highest levels in the anterior quadrant (Coffinier et al., 2001) (Fig. 6K), where anterior neural specification is taking place (Fig 6I). This pattern resembles the complementary expression of *Xenopus Chrd.1* and *nodal* genes in the BCNE/prospective brain and the NC (dorsal endoderm), respectively (Fig. 6A). However, it was not studied if Nodal normally controls mouse Chrdl1.

Remarkably, Nodal^-/-^ embryos showed an expanded and premature specification of neural ectoderm expressing forebrain markers in the epiblast (including the rostral forebrain regulator Hesx1), and Gsc expression persisted in the future GO region at pre-streak stages (Camus et al., 2006). When examined at streak-stages, Nodal^-/-^ embryos did not develop a morphologically distinguishable PS, and Gsc expression was always absent. Interestingly, Tbxt expression was also absent, except from some caudal cells in around 10% of mutant embryos. Moreover, some patches of other caudal mesoderm markers were present in 25% of mutant embryos (Conlon et al., 1994). FoxH1^-/-^ embryos neither expressed Gsc nor Foxa2 at the anterior tip of the PS during gastrulation and failed to form a definitive node and notochord (Yamamoto et al., 2001), (Hoodless et al., 2001). It is surprising that Chordin or Noggin expression were not analyzed in Nodal knockouts. However, conditional removal of Smad2 activity from the epiblast or deletion of the proximal epiblast enhancer (PEE) of the Nodal gene (which renders attenuated expression of Nodal in the PS) neither suppressed node formation nor anterior neural specification. However, Gsc expression corresponding to the prechordal plate was lost at the late-streak stage in both mutants, while Chordin and Noggin expression was aberrant and decreased along the axial midline in Smad2 mutants (it was not analyzed in Nodal-PEE mutants) (Vincent et al., 2003). Therefore, Gsc expression in the mouse AM region also appears to display two phases in relation to Nodal responsiveness, as those we show here for *Xenopus:* an earlier one (before PS formation) in the future GO region, which does not depend on Nodal (Camus et al., 2006), and a later one, in the node/prechordal plate, which requires Nodal (Conlon et al., 1994), (Yamamoto et al., 2001), (Hoodless et al., 2001), (Vincent et al., 2003). Overall, these results in mouse resemble our findings after blocking Nodal in *Xenopus,* indicating that this pathway is required for restricting forebrain induction before gastrulation (Camus et al., 2006), for the maintenance of PM cells during gastrulation and for the development of notochordal cells (Conlon et al., 1994), (Yamamoto et al., 2001), (Hoodless et al., 2001), (Vincent et al., 2003). However, development of a subset of posterior mesodermal cells does not show a high dependency on Nodal, both in mouse (Conlon et al., 1994) and *Xenopus* (this work). These observations suggest that in amphibians and mammals, before overt signs of gastrulation, Nodal is not necessary for the initial specification of the prechordal pre-organizer, but as gastrulation progresses, Nodal is required for the maintenance of the GO, the PM and the notochord.

In conclusion, while in *Xenopus,* the same gene *(Chrd.1)* is expressed in both pre-brain and pre-organizer territories in the BCNE (Fig. 6A), separate mouse genes encoding BMP antagonists belonging to the Chordin family are expressed in the presumptive neural plate (Chrdl1) and the GO (Chrd1) (Fig. 6K,L). In *Xenopus,* Nodal differentially regulates *Chrd.1* among the BCNE cells, being necessary for the initiation of *Chrd.1* expression in the presumptive CM sub-population and to restrict that of the pre-brain sub-population. In mouse, it is not known if Nodal analogously restricts Chrdl1 expression in the presumptive brain territory, although their complementary expression patterns suggest so. The SMAD2 conditional knockout indicates that Nodal is also necessary for Chrd1 expression during gastrulation in mouse, although it is not known if it is required for its initial expression in the GO or for its maintenance, since the analysis was performed at headfold stage (Vincent et al., 2003).

In chick embryos, the posterior marginal zone (PMZ) was proposed as the NC equivalent (Joubin and Stern, 2001) (Fig. 6G,H). The Köller’s sickle (KS) is a crescent shaped region of the superficial portion of the PMZ (Stern, 1990) (Fig. 6H). Two cell populations that contribute to the chick GO (Hensen’s node) were identified before the onset of gastrulation. One is initially located in the epiblast, just above the anterior face of KS at stage X and moves anteriorly between stages XI and XIII to the center of the blastoderm. These are known as central cells (CC) (Fig. 6H). The second group, known as posterior cells (PC), is located within the middle layer and is initially associated with the inner face of KS (Fig. 6H). These PC start expressing Gsc before gastrulation and migrate later than the CC, within the middle layer, together with the tip of the PS. When both populations meet at the mid-streak stage, they establish a completely functional GO (Izpisúa-Belmonte et al., 1993), (Streit et al., 2000), (Joubin and Stern, 2001). As in *Xenopus* and unlike in mouse, Chrd expression in chick embryos begins before the onset of gastrulation (Fig. 6G). Transcripts are initially found in a region just anterior to KS in the epiblast and in the underlying cells of the middle layer. Therefore, it is expressed in both populations that contribute to the GO (Streit et al., 1998) (Fig. 6G). A rough fate map of the early blastula/pre-streak chick embryo showed that the prospective forebrain position lies in the epiblast, also immediately anterior to the KS (Stern et al., 2006) (Fig. 6H), suggesting that Chrd is also transiently expressed in the chick forebrain precursors, like in the *Xenopus* BCNE at equivalent stages, as suggested by (Kuroda et al., 2004), (Reversade et al., 2005) (Fig. 6G). Ectopic expression of Chrd in the non-neural ectoderm of the area pellucida in gastrulating chick embryos was unable to induce a secondary PS or neural markers. However, when Chrd was ectopically expressed earlier, in the anterior edge of the area pellucida before the onset of gastrulation, a secondary axis with PS, node and neural ectoderm with the typical horseshoe shape of the anterior neural plate was induced (Streit et al., 1998). This suggests that forebrain precursors are competent to be recruited by Chrd before the onset of gastrulation in chick embryos.

In pre-streak chick embryos (stage XII/XIII), Nodal is expressed in a region confined to the middle two-thirds of KS, while Chrd expression is restricted to the midline region of the epiblast, just rostral to KS (Lawson et al., 2001) (Fig. 6G). Therefore, as in *Xenopus,* Nodal is expressed in the chick NC equivalent. Nodal alone was insufficient to induce ectopic Chrd in explants of anterior epiblast of pre-streak embryos but could induce it when combined with FGF8. On the other hand, in the absence of FGF8 signaling, Nodal expression remained unaffected in explants of posterior blastoderm (containing KS) of pre-streak embryos, while Chrd expression was decreased (Matsui et al., 2008). These results indicate that FGF8 signaling from the nascent hypoblast is necessary for Chrd expression before the onset of avian gastrulation and that Nodal might cooperate but is insufficient to induce it. However, experiments blocking Nodal were not performed to address if it is indeed required for triggering Chrd expression in the posterior epiblast before gastrulation.

In Zebrafish, *chrd* is readily detected well before gastrulation in the dorsal region, including the future GO region and extending more or less towards the animal pole, depending on the bibliography (Miller-Bertoglio et al., 1997), (Sidi et al., 2003), (Branam et al., 2010) (Fig. 6D). In contrast to the other vertebrate models discussed here, *chrd* is additionally expressed during gastrulation in territories beyond the GO, including the prospective brain and other neuroectodermal regions (Miller-Bertoglio et al., 1997), (Sidi et al., 2003), (Branam et al., 2010) (Fig. 6E,F). Notably, in double mutants for *nodal1* and *nodal2* (squint/cyclops) or in maternal/zygotic mutants for the essential cofactor for Nodal signaling *tdgf1 (MZoep* mutants), the *chrd* domain in the dorsal center of the zebrafish blastula was reduced in size, but strong expression persisted in a considerable subdomain (Gritsman et al., 1999). Also, at blastula stages, *tbxt* expression was suppressed in the dorsal region but persisted in the remaining presumptive mesodermal ring, while *gsc* expression was suppressed in these mutants (Feldman et al., 1998), (Gritsman et al., 1999). These observations indicate that the population of *chrd+* cells in the zebrafish blastula is also heterogeneous in Nodal requirements, as we found in *Xenopus* (this work). They also show that the CM precursors (as we found in *Xenopus)* as well as the PM precursors (unlike what we observed in *Xenopus)* require Nodal signaling for their initial specification (Fig. 6D). As both PM and CM markers were suppressed at blastula stages, the *chrd+* cells that remain in these mutants might as well represent future neural cells.

Interestingly, zebrafish has a *chordin-like2 (chrdl2)* gene which also behaves as a BMP antagonist. The *chrdl1* gene could not be identified in the genome (Branam et al., 2010). *Chrdl2* is strongly expressed throughout the animal hemisphere before gastrulation (Fig. 6D), thus introducing an additional level of BMP antagonism to the dorsal region. Knock-down experiments showed that *chrl2* is required together with *chrd* for dorsal development during the patterning of the embryonic dorsal-ventral axis (Branam et al., 2010).

### Concluding remarks

The rostral forebrain is an evolutionary acquisition of vertebrates related to the appearance of the Hesx1 gene at the beginning of vertebrate evolution (Ermakova et al., 1999), (Wilson and Houart, 2004), (Ermakova et al., 2007), (Bayramov et al., 2016) and derives entirely from the BCNE center (Kuroda et al., 2004). We have previously shown that expression of the rostral forebrain regulator *hesx1* persists after blocking Nodal signaling in *Xenopus* (Murgan et al., 2014). Indeed, the expansion of the Hesx1 domain in Nodal^-/-^ mouse embryos occurred before gastrulation (Camus et al., 2006). Altogether, our findings and the comparison between different vertebrate models indicate that the establishment of the CNS during the development of vertebrates requires not only a very early induction of the brain territory but also its delimitation. This process involves Nodal signaling in the differential segregation of the cell populations that give rise to the dorsal structures (brain and AM) as early as at the onset of neural induction at blastula stage.

## MATERIALS AND METHODS

### Embryological manipulations, mRNA synthesis and injections

Albino and wild-type *Xenopus laevis* embryos were obtained by natural mating or by in vitro fertilization using standard methods (Sive et al., 2010) and staged according to (Nieuwkoop and Faber, 1994). Parental animals were obtained from Nasco (Fort Atkinson, WI). Protocols were approved by the Laboratory Animal Welfare and Research Committee (CICUAL) from Facultad de Medicina-UBA.

To obtain synthetic capped mRNAs, the following plasmids were employed as templates: *Xcer-S pCS2+* (gift from Eddy de Robertis) (Bouwmeester et al., 1996), *pCS2 MT foxH1-SID* (gift from Uwe Strähle) (Chen et al., 1997), and *pCS2-NLS-lacZ* (gift from Tomas Pieler) (Bellefroid et al., 1996). These plasmids were linearized with NotI and transcribed with SP6 mRNA polymerase with the mMESSAGE mMACHINE SP6 Transcription Kit (ThermoFisher Scientific, AM1340), following the manufacturer’s instructions. Synthetic capped mRNAs were purified with the RNeasy Mini Kit (QIAGEN #74104).

Microinjection, culture and fixation of embryos were performed as previously described (Murgan et al., 2014). *Cer-S* mRNA was injected in the vegetal region of the four cells at s3 (0.5 ng/cell). *Foxh1-SID* mRNA (0.25 to 1 ng) was unilaterally injected at s2. Microinjections included as tracer 30-40 ng/cell of Dextran Oregon Green 488, MW 10000, anionic, lysine fixable (DOG; Thermo Fisher Scientific, D7171) or of Dextran Alexa Fluor 59, MW 10000, anionic, Fixable (Thermo Fisher Scientific, D22913); or 0.5 ng/cell of *nuc-lacZ* mRNA.

### *In situ* hybridization and X-gal staining

Plasmids for obtaining antisense RNA probes for whole mount *in situ* hybridization (ISH) were linearized and transcribed as follows. *Chrd1: pSB59-chrd1* (gift from Eddy de Robertis) (Sasai et al., 1994) was cut with EcoRI and transcribed with T7 RNA polymerase; *Gsc: gsc pG500* (gift from Ken Cho) (Cho et al., 1991), XbaI/T3; *myod1: pSP73-XmyoD* (gift from Cristof Niehrs) (Hopwood et al., 1989), BamHI/SP6; *not: pBS-KS-Xnot:* HindIII/T7 (gift from David Kimelman) (von Dassow et al., 1993); *sia1: pBluescript RN3 Xsia ORF* (gift from Patrick Lemaire) (Lemaire et al., 1995), HindIII/T7; *sox2: pBS sox2* (gift from Yoshiki Sasai) (Kishi et al., 2000), EcoRI/T7; *tbxt: apSP64T bra* (gift from Abraham Fainsod) (Smith et al., 1991), SalI/SP6. The preparation of digoxigenin-labeled antisense RNA probes and the whole-mount ISH procedure were performed as previously described (Pizard et al., 2004), except that the proteinase K step was omitted. X-gal staining for revealing the *nuc-lacZ* tracer was performed as previously described (Franco et al., 1999).

### RT-qPCR analysis

Embryos injected with *cer-S* mRNA and uninjected siblings were allowed to grow until stages 9 or 10. For total RNA extraction, 3 embryos at s9 or 8 embryos at s10 were placed in 1.5 ml tubes. After liquid withdrawal, tubes were placed on ice, and 200 μl (for s9 samples) or 400 μl (for s10 samples) of TRIreagent (Merck, Cat. No 93289) was added. Embryos were resuspended 10 times with micropipette. Samples were stored at −80°C until RNA extraction, which was performed following manufacturer’s instructions. 1 μg of RNA was treated with DNAse I (Ambion Cat. No.: AM2222) and used for first-strand cDNA synthesis, using High Capacity Reverse Transcription Kit with random primers (Applied Biosystems, Cat No. 4368814). Amplification was performed in triplicate in an Applied Biosystems 7500 Real-Time PCR System machine using Power Up SYBR Green Master Mix (Applied Biosystems, Cat. No. A25472). Melt curves were analyzed to confirm specificity of PCR product. Efficiency of each PCR amplification was estimated using the slope of a standard curve. Relative gene expression was calculated using Pfaffl’s mathematical model, with Histone H4 expression levels as standard.

The following primers were used for RT-qPCR: *Chrd.1* F: ACTGCCAGGACTGGATGGT, *Chrd.1* R: GGCAGGATTTAGAGTTGCTTC (Leibovich et al., 2018); *gsc F:* TTCACCGATGAACAACTGGA, *gsc R:* TTCCACTTTTGGGCATTTTC (Leibovich et al., 2018); *histone H4* F: GGCAAAGGAGGAAAAGGACTG, *histone H4* R: GGTGATGCCCTGGATGTTGT (Cao et al., 2007); *mix1* F: CAAAAGCCACCAAGCCCATT, *mix1* R: TGCTGAAGGAAACATTGCCC (Sun et al., 2015); *sox2* F: GAGGATGGACACTTATGCCCAC, *sox2* R: GGACATGCTGTAGGTAGGCGA E.M. De Robertis http://www.hhmi.ucla.edu/derobertis/.

### Data collection and statistics

*Cer-S* mRNA-injected batches for ISH analysis were tested by the effect on the mesodermal marker *myod1* at neurula stage and were used when *myod1* expression was strongly attenuated or abolished in injected embryos in comparison to uninjected siblings (Fig. 3A,B; Table 1). For RT-qPCR analysis, *cer-S* mRNA-injected batches were tested at s9 and s10 by the effect on the expression of *mix1*, a direct target of Nodal signaling (Charney et al., 2017), and were used when *mix1* expression was significantly reduced to less than 50% in relation to uninjected sibling controls (Suppl. Fig. 1).

Numbers of samples (n) and of biological replicates (N) analyzed are indicated for each set of experiments in the figures and tables. Biological replicates represent batches of embryos from independent mating pairs, or from different groups of embryos from the same batch in the case of RT-qPCR assays at s9. Statistical tests applied for RT-qPCR analysis are described in the corresponding Materials and Methods section and in the figures. Differences were considered significant when p<0.05.

## ACKNOWLEDGEMENTS

We thank the following colleagues for providing us with DNAs: Ken Cho (gsc), Eddy De Robertis *(Chrd.1, cer-S),* Abraham Fainsod *(tbxt),* Tomas Pieler (NLS-lacZ/*nuc-lacZ),* David Kimelman *(not),* Patrick Lemaire *(sia1),* Christof Niehrs *(myod1),* Yoshiki Sasai *(sox2),* and Uwe Strähle *(foxh1-SID). We* are grateful to Marianela Ceol Retamal, Andrea Pecile, Manuel Ponce, and Ezequiel Yamus for animal husbandry, and to María Belén Favarolo for help with experiments.

## COMPETING INTERESTS

The authors declare no competing or financial interests.

## AUTHOR CONTRIBUTIONS

Conceptualization: A.M.C.C., S.L.L; Formal analysis: A.M.C.C., M.B.T., L.E.B.L., S.L.L.; Investigation: A.M.C.C., M.B.T., L.E.B.L., S.L.L.; Drafting of original manuscript: A.M.C.C., S.L.L.; Writing – critical revision & editing: A.M.C.C., L.F.F, M.R., S.L.L.; Visualization: A.M.C.C., M.B.T., S.L.L.; Supervision: S.L.L.; Resources: S.L.L., M.R., L.F.F.; Project administration: S.L.L.; Funding acquisition: S.L.L.

## FUNDING

Research was supported by Agencia Nacional de Promoción Científica y Tecnológica, Argentina (PICT 2011-1559, PICT 2014-2020) and Consejo Nacional de Investigaciones Científicas y Técnicas, Argentina (PIP 2012-0508, PIP 2015-0577; fellowship to A.M.C.C.).

## Notes

### Competing Interest Statement

The authors have declared no competing interest.

