## Supplemental Figure 1 for "Segregation of brain and organizer precursors is differentially regulated by Nodal signaling at blastula stage"

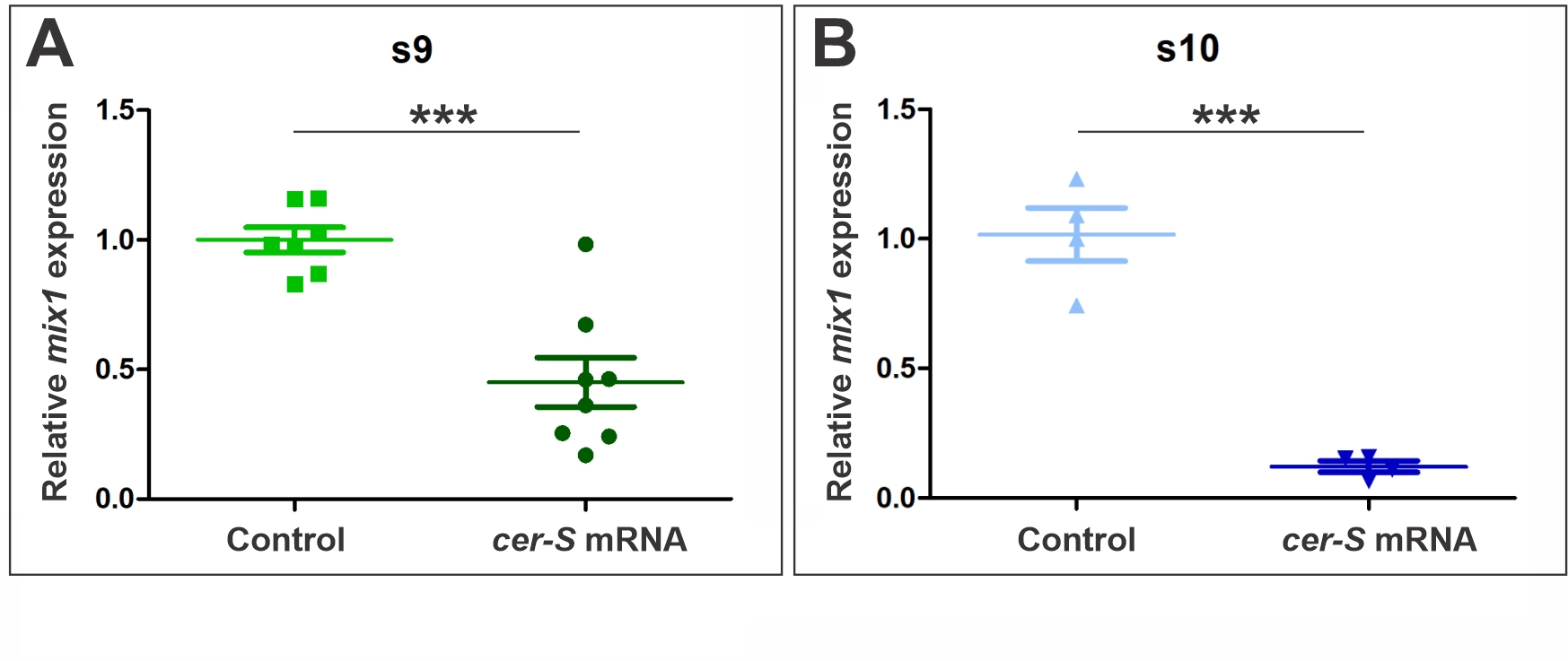

**Supplementary Figure 1.** Relative expression levels of *mix1* transcripts at s9 (A) and s10 (B) in *cer-S*-injected embryos and uninjected sibling controls, analyzed by RT-qPCR. Values for each biological replicate are indicated by symbols. *Cer-S*-injected biological replicates were tested by the effect on the expression of *mix1*, a direct target of Nodal signaling (Charney et al., 2017). *Mix1* expression was significantly reduced (p<0.05, unpaired, two-tailed t-test) in *cer-S-*injected embryos in relation to uninjected sibling controls, both at s9 (p=0.0003) and at s10 (p=0.0001). Only those biological replicates that showed *mix1* expression reduced to less than 50% in relation to uninjected sibling controls were used for RT-qPCR analysis of the other markers shown in this work.

**Charney, R. M., Paraiso, K. D., Blitz, I. L. and Cho, K. W. Y.** (2017). A gene regulatory program controlling early Xenopus mesendoderm formation: Network conservation and motifs. *Semin. Cell Dev. Biol.* **66**, 12–24.
